# The effect of intercellular medium salinity on the dielectric constant of cell membranes and the excitation thresholds of the action potential

**DOI:** 10.1101/2023.07.24.550397

**Authors:** M. N. Shneider, M. Pekker

## Abstract

A mechanism for axon swelling and subsequent compression in the area of action potential passage is considered. It is shown that the dynamics of changes in the radius of the axon during the passage of the action potential can be explained by changes in osmotic pressure. Axon swelling occurs at the leading edge of the action potential when Na^+^ ions flow from the extracellular electrolyte into the intracellular space, and subsequent compression occurs at the trailing edge during the ejection of K^+^ ions. The change in dielectric constant of the axon membrane is also discussed, and a possible explanation for the decrease in the excitation threshold of the action potential with an increase in salinity of the external electrolyte is proposed.

## I. Introduction

In [1−6] it was shown that during the passage of an action potential in nerve fibers, for example, in the axons of a squid, the radius of the axons follows the action potential, that is, it initially increases and then decreases. The authors attribute this effect to the injection of sodium ions into the axon during the passage of the action potential. The conclusions drawn in [5, 6] indicate that the swelling of the squid axon in the region of action potential cannot be explained by the redistribution of fluid inside the axon, which occurs during mechanical excitation of surface waves.

Numerous works have been devoted to the experimental study of the effect of the ionic composition of electrolyte on nerve fibers. In this paper, we will highlight only a few of these works, those which—in our opinion—are the most important. It was shown in [7] that an increase in the amount of divalent ions in the external electrolyte reduces the potential difference between the internal and external electrolytes in the squid axon, significantly lowering the threshold of sodium channel opening. Based on the analysis of X-ray diffraction data, [8] reported that high concentrations of KBr and KCl salts lead to an increase in the distance between phospholipid heads and, accordingly, to an almost twofold decrease in the van der Waals forces of attraction between them.

During the injection of sodium ions into the area of the action potential passage, the local salinity of both the internal electrolyte and external electrolyte increases. As a result of osmosis, previous experiments [1– 6] indicate an outflow of water from the axon, leading to its compression rather than swelling. The reasons for axon swelling at the growth stage of the action potential are discussed in Part II of our work. This section proposes that the additional osmotic pressure drop due to the redistribution of ions between the outer and inner sides of the membrane determines the dynamics of axon radius change during the duration of an action potential.

In [7], it was shown that the addition of divalent chloride salts such as CaCl_2_, MgCl_2_, and MnCl_2_ to a solution of NaCl_2_ reduces the potential difference (resting potential) across the membrane of nerve fibers, although it would seem that the effect should have been the opposite. Indeed, an increase in the salinity of the solution leads to an increase in the surface charge on the membrane and, accordingly, to an increase in the potential difference on 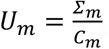, where *Σ*_*m*_ is the surface charge density, *C*_*m*_= *ε*_*m*_*ε*_0_/*d*_*m*_ is the membrane capacitance per unit area, *ε*_*m*_ is the membrane dielectric constant, *ε*_0_ is the vacuum permittivity, and *d*_*m*_ is the membrane thickness.

The experiments carried out in [8] showed that an increase in electrolyte salinity leads to an increase in membrane porosity, that is, an increase in the distance between phospholipid heads. Therefore, it can be expected that an increase in the density of ions in the electrolyte will also increase the amount of water in the membrane. In Part III of this work, saturation of the membrane with water is found to increase the dielectric constant of the phospholipid membrane, which leads to a decrease in the potential difference on the membrane of nerve fibers and, accordingly, lowers the opening threshold of sodium channels. Since water within the phospholipid membrane significantly influences the effective dielectric constant, an increase in the amount of water in the membrane is expected to result in a corresponding increase in the dielectric constant. In our opinion, saturation of the membrane with water explains the decrease in resting potential in the nervous tissue observed in [7] and, accordingly, the decrease in the threshold for opening transmembrane sodium channels.

### II. Osmotic pressure acting on a bilayer phospholipid membrane in an electrolyte

It was found experimentally [1, 2] that at the leading edge of the action potential, the time dependence of the change in the radius of the squid axon exactly follows the dynamics of the action potential and reaches ∼1 nm at the moment when the action potential reaches its maximum. In the recovery phase, when the action potential decreases, the radius of the axon also decreases and, at the action potential minimum point, becoming 1–4 nm less than the equilibrium value. Then it slowly recovers.

Let us evaluate the role of changes in the osmotic pressure on the membrane due to a local increase in the density of sodium ions in the region of action potential. Typical intracellular and extracellular ion densities are shown in Table 1 [9].

**Table 1.** Intracellular and extracellular ion densities (following [9]).

| Ions | Squid Axon |  | Muscle Fibers (neurons) |  |
| --- | --- | --- | --- | --- |
|  | Intercellular mM/L | extracellular mM/L | Intercellular mM/L | extracellular mM/L |
| $\text{Na}^+$ | 50 | 460 | 10 | 125 |
| $\text{K}^+$ | 400 | 10 | 124 | 2 |
| $\text{Ca}^{2++}$ | — | 10 | 5 | 2 |
| $\text{Mn}^{2++}$ | 10 | 54 | 14 | 1 |
| total | 460 | 534 | 153 | 130 |
| $\text{Cl}^-$ | 135 | 560 | 2 | 77 |
| $\text{HCO}_3^-$ | — | — | 12 | 27 |
| (A)<sup>-</sup> | 345 | — | 74 | 13 |
| Others | — | — | 65 | 13 |
| Total | 460 | 560 | 153 | 130 |

Let the density of sodium ions near the inner surface of the membrane increase by 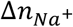. In this case, the excess osmotic pressure on the membrane surface increases as follows [10]:

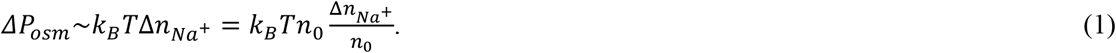

A similar estimate can be written for the change in osmotic pressure drop as a result of the outflow of K^+^ ions at the trailing edge of the action potential.

The local release of potassium ions from the axon to the outside at the stage of the fall of the action potential stimulates the reverse process: the outflow of water from the axon to the outside. Both these processes are accompanied by a local increase in excess osmotic pressure, first on the inner side of the membrane, and then on the outer side. In our opinion, this explains the results of experiments [1−6] - an increase in the radius of the axon at the initial stage of the action potential and its subsequent decrease at the final stage.

### III. Change in the dielectric constant of the membrane with increasing salinity of the solution

Let us briefly describe the setting of experiments and the results of [7], which showed that the addition of divalent chloride salts such as CaCl_2_, MgCl_2_, and MnCl_2_ to a solution of NaCl reduces the voltage across the membranes nerve fibers. In [7], divalent chloride salts were added to a 120 mM/L NaCl solution in a vessel with frog myelinated nerve fibers, and the potential difference between the inner and outer surfaces of the membrane was measured. The experiments show that adding 0.1 mM/L CaCl_2_ results in the measured potential difference *U*_*m*_ = −115 mV, adding 2 mM/L leads to *U*_*m*_ =−90 mV, and adding 6.0 mM/L leads to *U*_*m*_ = −80 mV. Similar results were obtained with the addition of other divalent chloride salts (e.g., ZnCl_2_). Thus, adding 0.07 mM/L results in *U*_*m*_ = −100 mV, adding 0.2 mM/L results in *U*_*m*_ = −95 mV, and adding 2 mM/L results in *U*_*m*_ = −75 mV.

On the other hand, it was shown in experiments [8] that an increase in electrolyte salinity leads to an increase in membrane porosity, that is, an increase in the distance between phospholipid heads. Therefore, it can be expected that with an increase in salinity, which is unambiguously associated with an increase in the density of ions in the electrolyte, the amount of water in the membrane also increases. The water contained in the phospholipid membrane contributes to the effective permittivity. Therefore, an increase in the amount of water in the membrane should lead to an increase in its dielectric constant.

Considering the water contained in the lipid membrane as an impurity with a relative volume, to estimate the effective permittivity of the membrane *ε*_*m*_, it is convenient to use the simplest approximate Rayleigh formula for the permittivity of two substances, one of which is continuous, and the second is uniformly distributed small droplets of the impurity substance [11]:

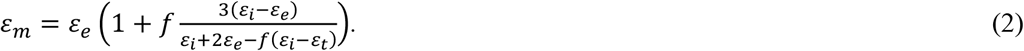

Here, *ε*_*m*_, *ε*_*e*_, and *ε*_*i*_ are the respective permittivities of the mixture, the main medium, and the impurity droplets. From (2), it follows that at *f* → 0: *ε*_*m*_ → *ε*_*e*_, and at *f* → 1: *ε*_*m*_ → *ε*_*i*_. Substituting in (2) the values of the permittivity *ε*_*e*_ = 2 for a dehydrated phospholipid membrane [12–14] and *ε*_*i*_ = *ε*_*w*_ 81 for water, we obtain the dependence of the membrane permittivity *ε*_*m*_ according to the volume fraction of water *f* (see Fig. 1).

**Figure 1.**
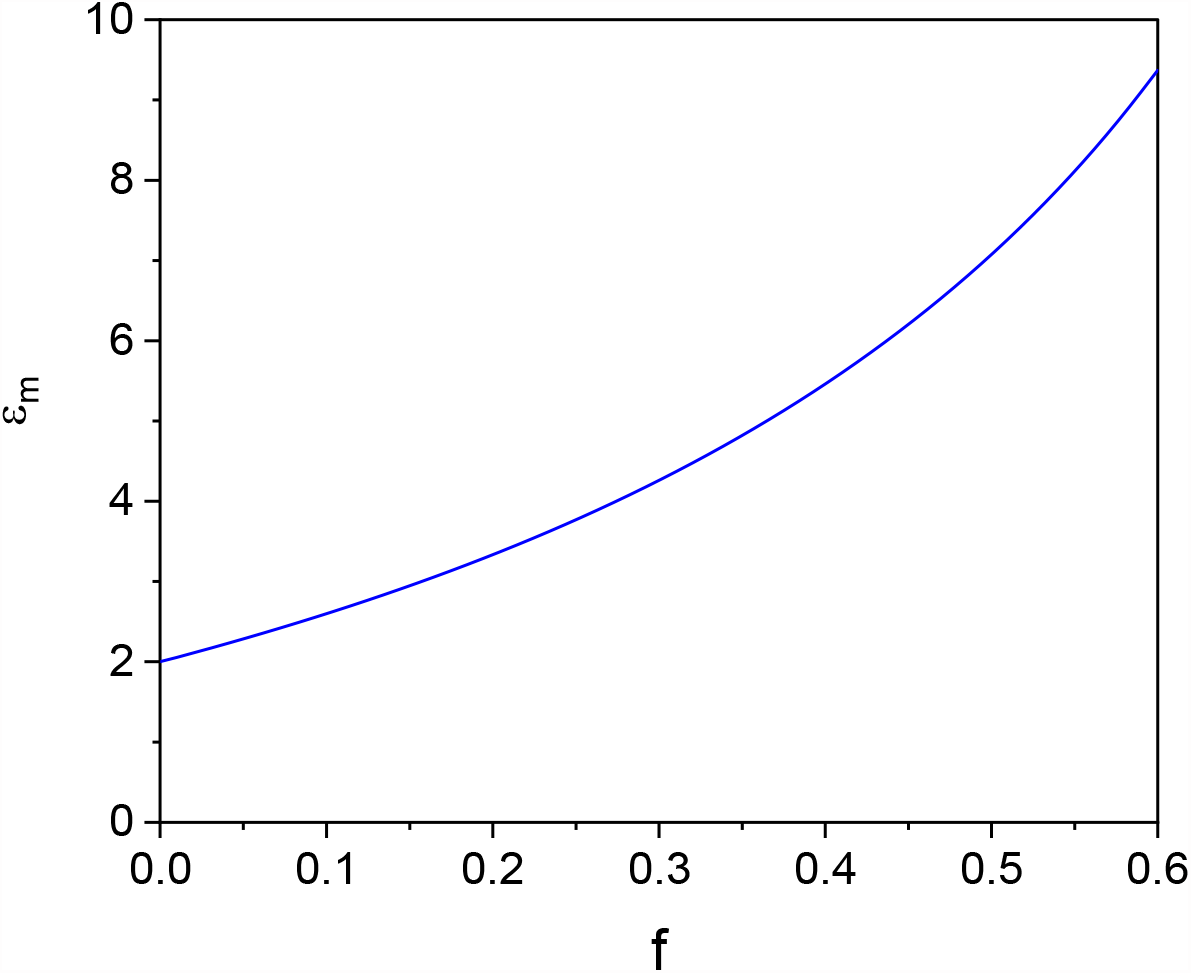
Dependence of the effective dielectric constant of the membrane *ε*_*m*_ on the water content *f* for *ε*_*e*_ 2 and *ε*_*i*_ = *ε*_*w*_ = 81, calculated by Formula (2)

Since the capacitance of a unit area of the membrane is equal to *C*_*m*_ = *ε*_*m*_ *ε*_0_/*d*_*m*_, then the potential difference across the membrane with the surface charge density *σ*_*m*_ is equal to

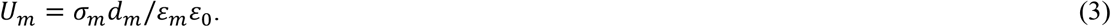

If we assume that with an increase in water salinity, the surface charge on the membrane and its thickness increase slightly compared to the water content, then *U*_*m*_∼1/*ε*_*m*_. The water content dependence of the ratio *U*_*m*_(*f*)/*U*_*m*_(*f* = 0) is shown in Fig. 2.

**Figure 2.**
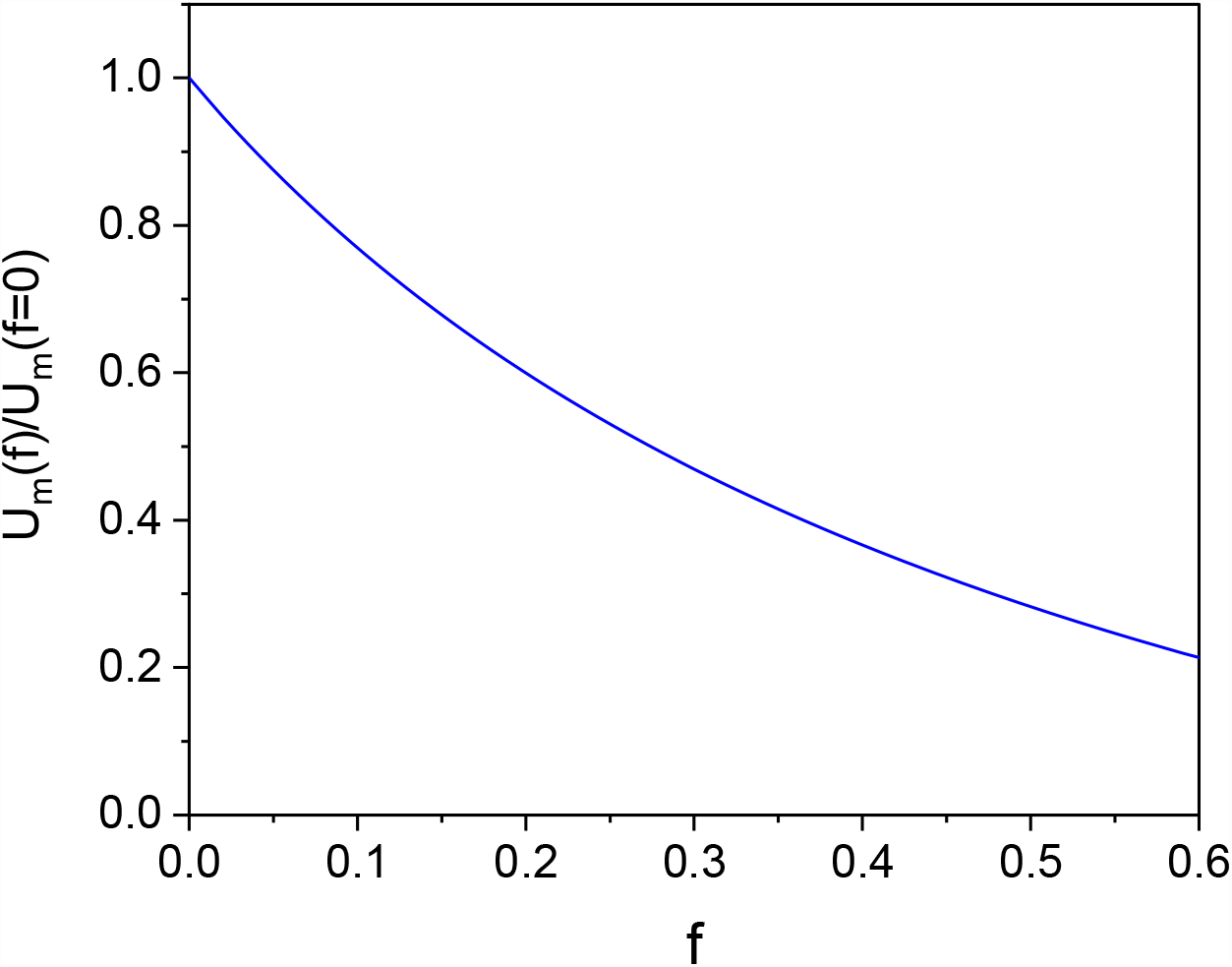
Dependence of the potential difference across the membrane on the water content *f* at a fixed value of the surface charge *σ*_*m*_ and thickness *d*_*m*_.

In our opinion, the saturation of the membrane with water explains the marked decrease in resting potential in the nervous tissue observed in [7] and, accordingly, the decrease in the threshold of sodium channel opening (threshold of action potential excitation).

Note that the considered effect of the dependence of potential difference across the membrane on water content at the same value of the surface charge can be directly related to the selective effect of non-equilibrium low-temperature plasma interacting with various cell cultures in a Petri dish (e.g., healthy and cancerous cells) observed in many experiments (see, for example, [15–18]). As was shown in [19], the interaction of non-equilibrium plasma with a physiological solution changes the ionic composition of the solution and, consequently, the drop of osmotic pressure across the cell membranes. In [19], it was supposed that the displacement of water from membranes and, consequently, the change in their dielectric constant, are different in healthy and cancer cells due to a noticeable difference in the mechanical properties of these cells. If, in addition, electric currents are induced in the solution, charging the membranes of these cells, then with the same additional charge on the membranes, the voltage on the membrane and the probability of its electroporation will be different for different cells. This, as was stated earlier in [20], may be one of the reasons for the observed selective effect of non-equilibrium plasma on healthy and diseased cells in a physiological solution in a Petri dish.

## IV. Conclusions

Changes in osmotic pressure associated with the flow of sodium ions from the external to the internal electrolyte at the stage of the growth of the action potential and the outflow of potassium ions at the stage of the fall of the action potential explain the dynamics of the axon radius observed in [1−6].

An increase in water content in the phospholipid membrane increases the dielectric constant of the membrane, which, at the same surface charge density, leads to a decrease in the potential difference across the nerve fiber membrane and, accordingly, lowers the action potential initiation threshold found in [7].

## Acknowledgments

The authors are grateful to Xuewei Zhang for valuable discussions. M.N.S. acknowledges partial support by the Princeton Collaborative Research Facility (PCRF) supported by the U.S. DOE under contract No. DE-AC02-09CH11466.

## Data Access Statement

All relevant data are within the paper.

